# Isolation and Characterisation of Hair Follicle-Derived Melanocytes

**DOI:** 10.64898/2026.01.14.699599

**Authors:** Fuyue Wu, Qingmei Liu, Xiaomin Shao, Shujun Chen, Jinran Lin, Jing Chen, Jian Li, Qi Zhang, Zheng Lin Tan, Wenyu Wu

**Affiliations:** ReMed Regenerative Medicine Clinical Application Institute, Shanghai, People’s Republic of China; Department of Dermatology, Huashan Hospital of Fudan University, Shanghai, People’s Republic of China

**Keywords:** melanocyte, melanocyte stem cell, melanocyte progenitor cell, hair follicle, NB-UVB

## Abstract

Melanocytes are unique population of cells which produce melanin pigment. However, the property of melanocytes from hair follicle and epidermis, particularly, their molecular signatures remained largely unknown. In this study, we investigated the properties of epidermis and hair follicle melanocytes by biochemical and molecular biological assays. We found that epidermal melanocytes are differentiated and mature form of hair follicle melanocytes, and they are potentially related. Irradiation with NB-UVB did not induce cell death of hair follicle melanocytes as observed in epidermal melanocytes, but activate hair follicle melanocytes, promote proliferation and migration. In situ study suggested that after UVB irradiation, root sheath of hair follicles, particularly proximity of interfollicular epidermis and hair bulb become darker, suggesting migration and maturation of hair follicle melanocytes after UVB irradiation, rather than cell death. This observation provides hints and evidence to the relationship between hair follicle and epidermal melanocytes, helps to explain the contradiction of melanocytes cell death after UVB irradiation and the use of UVB in treatment of vitiligo lesion, as well as the perifollicular repigmentation phenomenon observed in vitiligo lesion after NB-UVB phototherapy.

---

Melanocytes are unique populations of cells which synthesise melanin pigment. Melanin pigment is encapsulated in specialised organelles, known as melanosome (Bastonini et al., 2016). Melanin is transferred to keratinocytes, protecting hair follicles and skin from ultraviolet radiation-induced damage.

Unlike mice, studies have discovered melanocyte stem/progenitor cells present in bulge of hair follicles (Horikawa et al., 1996; Narisawa et al., 1997), sweat glands (Nakamura et al., 2015), adipose tissue (Ikeda et al., 2021) and basal membrane of epidermis (Lin and Fisher, 2007) in human. These melanocyte stem/progenitor cells are generally amelanotic, and they expressed different antigenic properties (Tobin & Bystryn, 1996). Comparative studies of epidermal and hair follicle melanocytes have revealed 3 morphological and antigenic distinct populations of melanocytes (Tobin & Bystryn, 1996). However, prior studies have not uncovered the molecular differences of these melanocytes.

While epidermal melanocytes are generally maintained by epidermal melanocyte stem/progenitor cells, it has been observed in mice that quiescent hair follicle melanocyte stem cells can be activated by UV exposure, and migrate to dermis and epidermis (Moon et al., 2017). While some studies have inferred the phenomenon in human by observing perifollicular repigmenetation of vitiligo after narrow band-UVB (NB-UVB) phototherapy (Nishimura et al., 2010; Quan et al., 2004), and perspective on human hair bulge amelanotic melanocytes are precursor cells of melanocyte has been described (Nishimura, 2011), the relationship between hair follicle melanocyte and epidermal melanocyte remained largely unknown.

In addition, effect of UVB irradiation on hair follicle melanocyte remained uncertain. Studies have also shown that UVB irradiation can induce cell death in epidermal melanocyte (Xu et al., 2025). If UVB irradiation has induced melanocyte cell death, it is unlikely that these cells will proliferate migrate to epidermis.

In this article, we sought to characterise the melanocyte in hair follicle in more detail. Our main goal is to attempt to explain what define human hair follicle melanocyte precursor, and what happen to hair follicle melanocyte after UVB irradiation.

## Results

### Identification and pseudotime trajectory analysis of epidermal and hair follicle melanocytes

While prior studies have reported the potential of hair follicle melanocyte as the stem cell reservoir of epidermal melanocytes, particularly through perifollicular repigmentation of vitiligo lesion after narrow band UVB phototherapy and inflammation (Qiu et al., 2015; Yamada et al., 2013; Inomata et al., 2009). However, neither direct observation nor inference based on available evidence based on melanocyte development has been reported. Therefore, it is important to first characterise both epidermal and hair follicle melanocytes isolated from their environment.

A distinct cluster of melanocytes can be identified based from uniform manifold approximation and projection (UMAP) mapping of data obtained from single-cell ribonucleic acid sequencing (scRNA-seq) (Figure 1a). This cluster is characterised by expression of melan-A (MLANA), pre-melanosome protein (PMEL), and microphthalmia-associated transcription factor (MITF) (Figure 1b), which are generally considered the markers of melanocytes.

**Figure 1.**
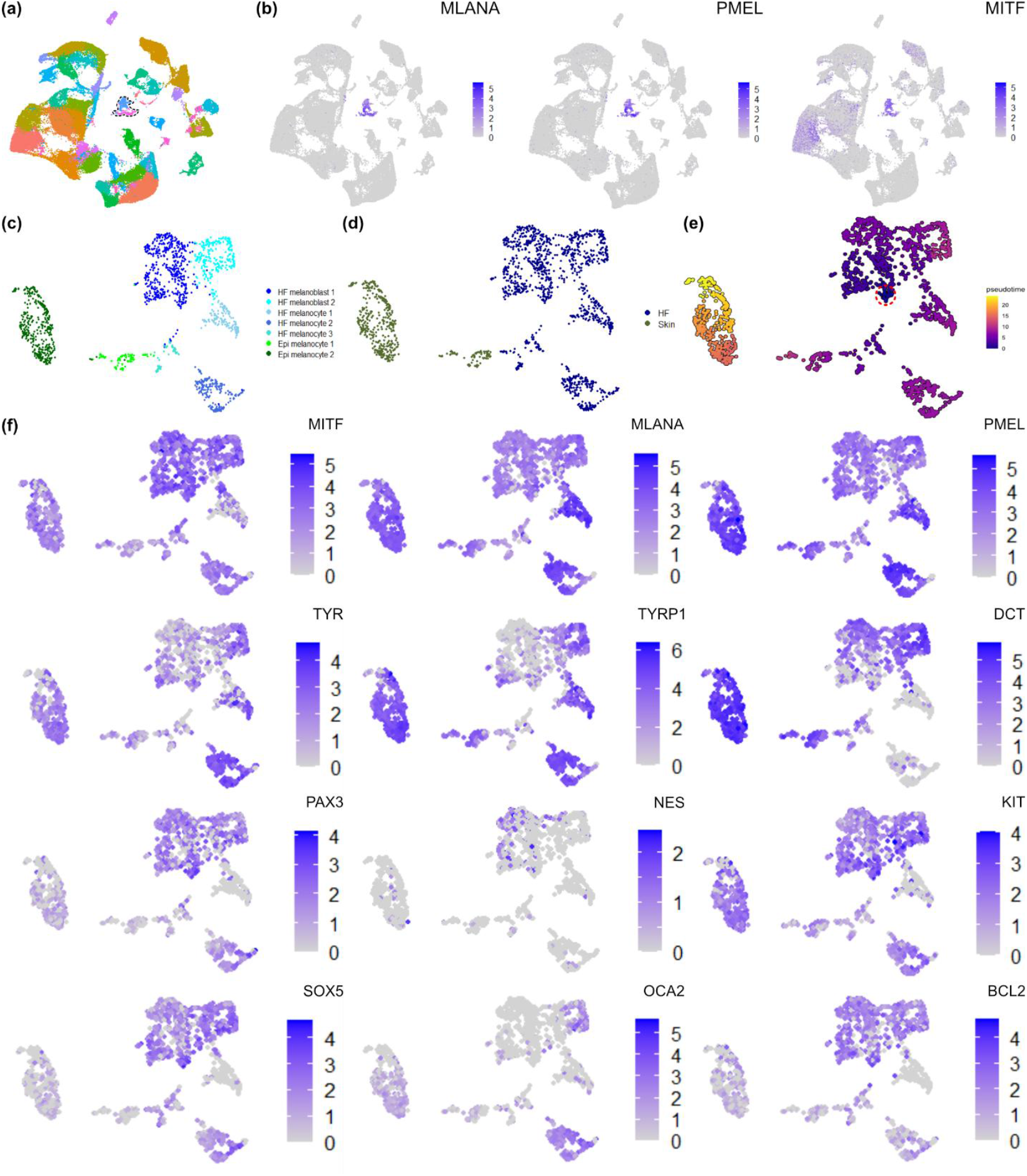
Comparison of molecular signature of skin and hair follicle melanocytes based on scRNA-seq. (a) Uniform manifold approximation and projection mapping of scRNA-seq data from skin and hair follicles. Clusters enclosed with dashed line are clusters representing melanocytes. (b) Mapping of general markers for melanocytes. (c) Uniform manifold approximation and projection mapping of melanocytes subset from this data and annotation for each cluster. HF: hair follicle, Epi: epidermal. (d) Ground truth of the origins of melanocytes. (e) Pseudo-time analysis of melanocytes. The region enclosed with dashed line is the origin of pseudo-time trajectories analysis. (f) Expression of genes related to epidermal and hair follicle melanocytes.

After isolation of the cluster representing melanocytes, dimensional reduction was conducted, and each cluster was annotated. Various clusters were identified, which includes amelanotic melanoblast (HF melanoblast 1), melanotic melanoblast (HF melanoblast 2), sacrificial melanocytes which probably representing the cluster which fail to dedifferentiate into melanoblast (HF melanocyte 1), bulb melanocyte (HF melanocyte 2), and transitioning melanocytes which will differentiate into epidermal melanocytes (HF melanocyte 3) from hair follicle (HF); interfollicular epidermal melanocytes (Epi melanocyte 1) and epidermal melanocyte (Epi melanocyte 2) from epidermis (Figure 1c) were annotated. Comparing the annotation to ground truth (Figure 1d) which represents the source of melanocyte suggested that the annotation well-represented the origin of cells, and epidermal melanocytes (primarily from interfollicular epidermis) can also be detected in hair follicles. The signature genes of each subcluster were shown in Figure 1f.

Pseudo-time trajectory analysis based on UMAP mapping of melanocytes suggested that melanoblast can undergo maturation, differentiate into bulb melanocytes, and into epidermal melanocytes through different trajectories (Figure 1e).

Pseudo-time trajectory analysis which is based on similarity and gene expression of cells, might be useful to infer progression of biological processes of cells. While it is difficult to observe differentiation of human hair follicle melanocyte into human epidermal melanocyte *in vivo*, result obtained from pseudo-time trajectories analysis based on markers related to stemness of melanocytes as origin might provide indirect evidence of the evolution of melanocytes.

### Characterisation of hair follicle melanocytes

To investigate the difference between hair follicle melanocyte and epidermal melanocyte, melanocytes were isolated and expanded from epidermis and hair follicle. Dermal fibroblast which is melanin nonproducing bipolar cell was used as control. Figure 2a shows that hair follicle melanocytes exhibit different morphology compared to epidermal fibroblast.

**Figure 2.**
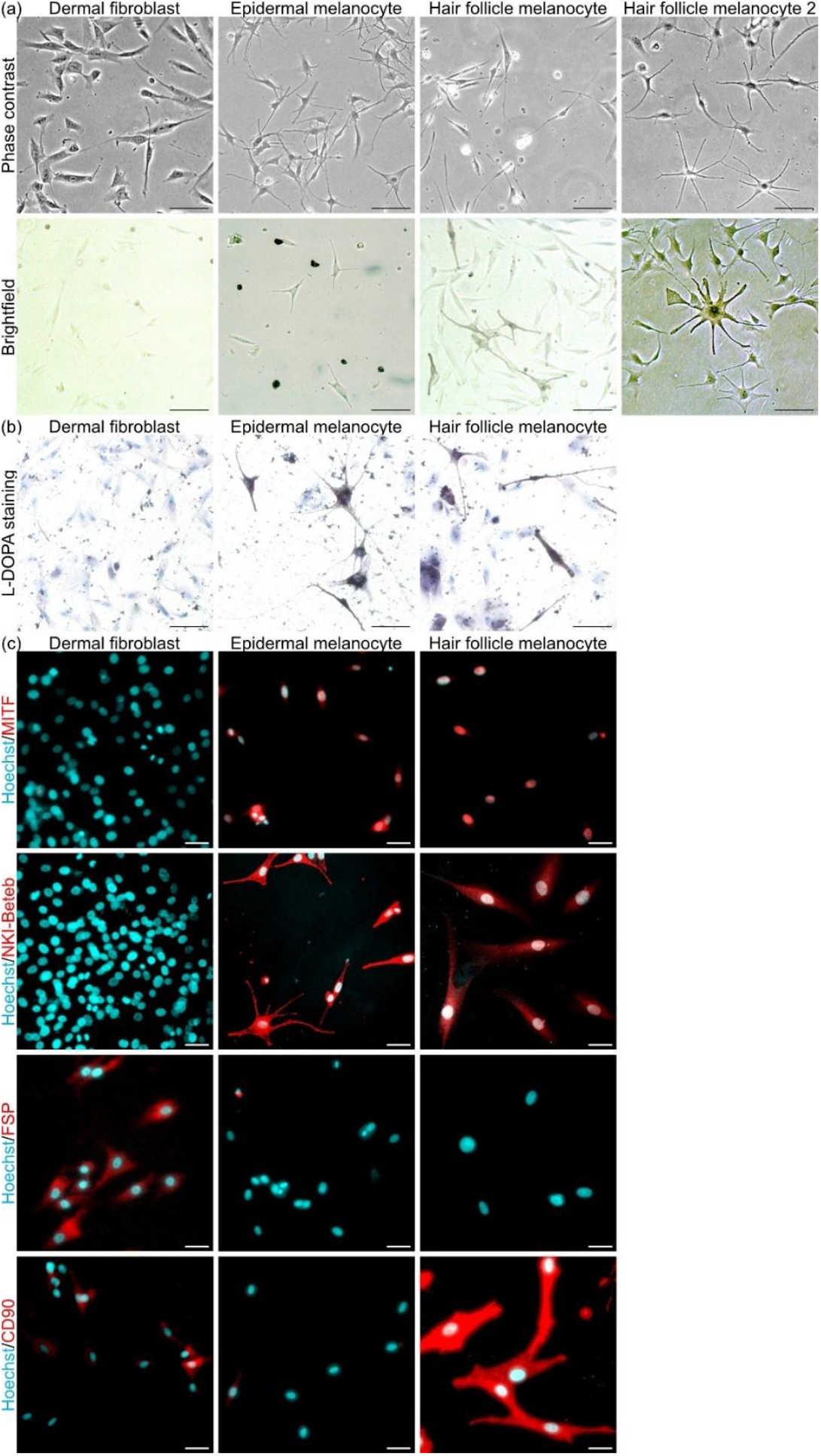
Characteristics of hair follicle melanocytes. (a) Micrographs of dermal fibroblast, epidermal melanocytes and hair follicle melanocytes. The scale bars are 100 μm. (b) *L*-DOPA staining of dermal fibroblasts, epidermal melanocytes and hair follicle melanocytes. Nuclei were counter-stained with haematoxylin. The scale bars are 50 μm. (c) Staining of dermal fibroblasts, epidermal melanocytes and hair follicle melanocytes with markers of melanocytes and fibroblasts. The scale bars are 20 μm.

While both dermal fibroblast and hair follicle melanocyte are bipolar, hair follicle melanocytes are smaller in size, and they are pigmented under brightfield microscope. Another type of melanocytes isolated from hair follicle exhibit dendrite morphology, densely pigmented, and their size are larger than epidermal melanocytes. However, this subpopulation did not proliferate in our culture medium.

The result of *L*-DOPA staining shows that a portion of epidermal and hair follicle melanocytes are *L-*DOPA positive cells (Figure 2b), which indicates that these cells are capable of melanin synthesis.

Further investigation was conducted via immunocytochemical staining. All epidermal and hair follicle melanocytes are MITF^+^ and NKI/beteb^+^ cells, which indicates that regardless of their melanin synthesis capability defined by *L*-DOPA staining, they are positive of melanocytes markers; while fibroblast surface protein (FSP) was used as negative markers for the bipolar morphology of hair follicle melanocyte resemble the morphology of mesenchymal cells. CD90 was also stained to confirm the stemness of hair follicle melanocyte, and the result suggested that cultured hair follicle melanocytes are CD90^hi^ cells compared to CD90-epidermal melanocytes and CD90^lo^ dermal fibroblast.

### Transcriptomic profile of hair follicle melanocytes

To further characterise hair follicle melanocytes, transcriptomic analysis was conducted on cultured hair follicle melanocyte, epidermal melanocyte and dermal fibroblast. Clustering by principal components (Figure 3a) suggested that technical variation among different group of samples was small, and these cells exhibit different transcriptomic property. Sample clustering analysis (Figure 3b) suggested that hair follicle melanocyte share the property of both dermal fibroblast and epidermal melanocyte, with higher similarity to dermal fibroblast. Comparing the up-regulated (Figure 3c) and down-regulated (Figure 3d) genes of hair follicle and epidermal melanocyte relative to dermal fibroblast shows that 1,185 genes can be used to characterise melanocytes as common genes, with 3,410 genes define epidermal melanocyte, and 559 genes define hair follicle melanocyte.

**Figure 3.**
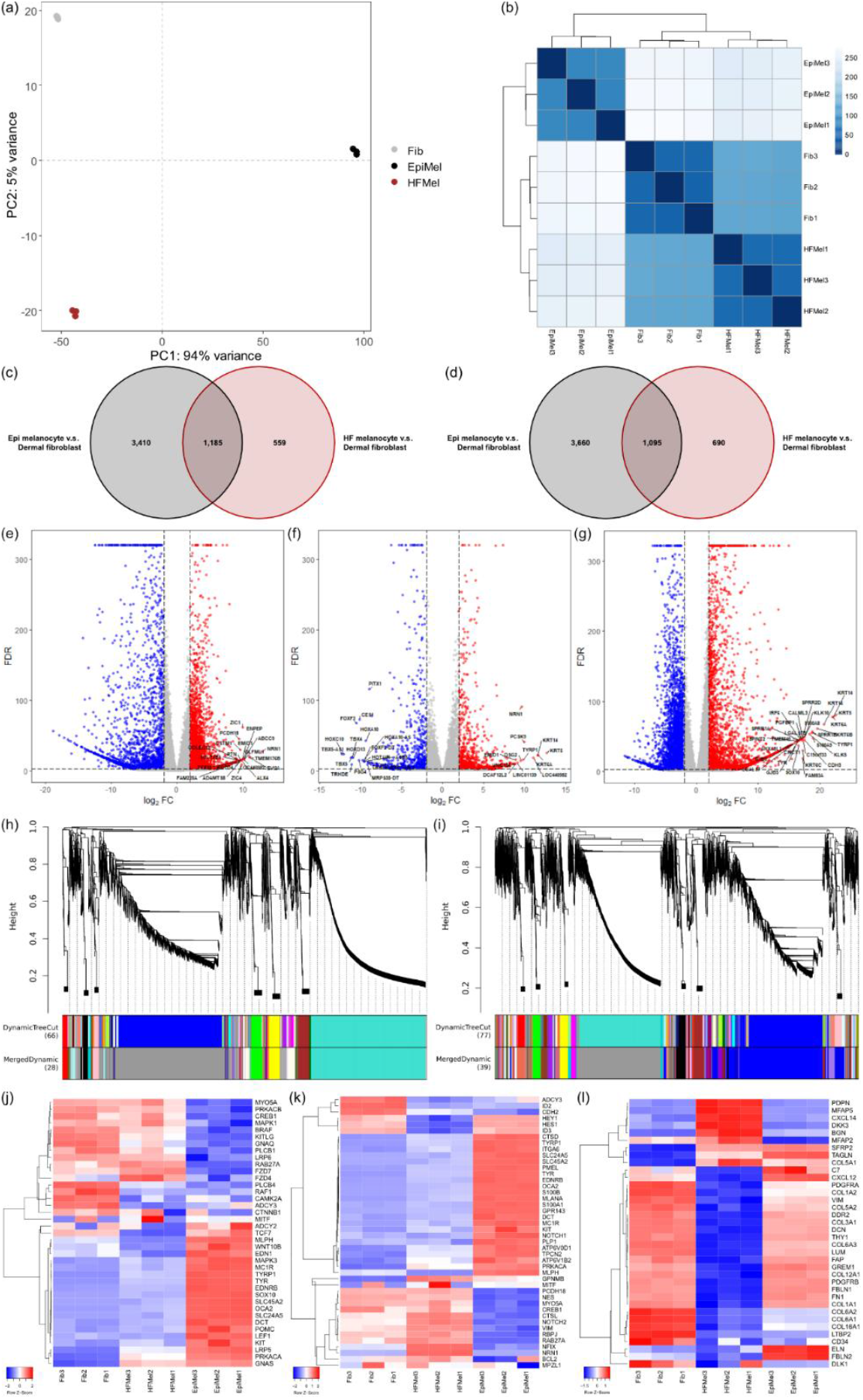
Transcriptomic profile of hair follicle melanocyte compared to epidermal melanocyte and dermal fibroblast. (a) Principal components analysis of hair follicle melanocyte, epidermal melanocyte and dermal fibroblast. Fib: fibroblast, EpiMel: epidermal melanocyte, HFMel: hair follicle melanocyte. (b) Clustering analysis among hair follicle melanocyte, epidermal melanocyte and dermal fibroblast. Venn diagram of (c) up-regulated and (d) down-regulated genes of epidermal melanocyte and hair follicle melanocyte in relative to dermal fibroblast. Volcano plot (e) hair follicle melanocyte in relative to epidermal melanocyte, (f) hair follicle melanocyte in relative to fibroblast and (g) epidermal melanocyte in relative to fibroblast. Cluster dendrogram obtained from WGCNA comparing (h) hair follicle melanocyte and epidermal melanocyte, and (i) hair follicle melanocyte and dermal fibroblast. Heatmap of genes related to (j) melanogenesis, (k) stemness and differentiation of melanocyte, (l) markers of dermal fibroblast.

Differential expression genes analysis of hair follicle melanocyte relative to epidermal melanocyte (Figure 3e) suggested that hair follicle melanocyte can be characterised by its expression of neuronal characteristics, e.g., neuritin 1 and protocadherin 18. As it is generally regarded melanocytes were derived from neural crest stem cells (Wang et al., 2024; Mort et al., 2015), expression of neuronal markers suggested that hair follicle melanocytes are in their early stages of development compared to epidermal melanocyte. On the other hand, comparing hair follicle melanocyte to dermal fibroblast (Figure 3f) enriched melanogenesis-related genes in hair follicle melanocytes, e.g., tyrosinase related protein 1, which characterise hair follicle melanocyte as a subpopulation of melanocyte. Comparing epidermal melanocyte to dermal fibroblast has also enriched genes related to melanogenesis, e.g., tyrosinase and tyrosinase related protein 1.

To further characterise hair follicle melanocyte, weighted gene co-expression network analysis (WGCNA) was conducted to identify identity biomarkers. While the sample size was small, and the result is subjected to noise and variation, it is sufficient to provide hint to the characteristics of hair follicle melanocyte. Cluster dendrogram of WGCNA of hair follicle melanocyte and epidermal melanocyte (Figure 3h) identify grey and turquoise module are highly correlated among samples. Turquoise module which enriched 4,956 genes characterising epidermal melanocytes, has a correlation coefficient of 1.00 and *p* < 0.00001. Analysing these genes suggested that they are related to cellular differentiation, suggesting that epidermal melanocyte is in differentiated states compared to hair follicle melanocyte; cluster dendrogram of WGCNA of hair follicle melanocyte and dermal fibroblast (Figure 3i) identify grey and blue module are highly correlated among samples. Grey module which enriched 2,556 genes characterising hair follicle melanocytes, has a correlation coefficient of 1.00 and *p* < 0.000001. Analysing these genes suggested that they are related to pigment metabolism, which include pigment biosynthesis and melanin biosynthesis, suggesting that hair follicle melanocytes are melanin synthesis cells, which the results were in align with differential expression genes analysis.

Heatmap plot suggested that while hair follicle melanocyte express melanogenesis-related genes (Figure 3j), the expression level is lower than epidermal fibroblast. Genes related to differentiation of melanocyte expressed by hair follicle melanocyte was lower than epidermal melanocyte, while expression of genes related to stemness were higher in hair follicle melanocyte (Figure 3k). Meanwhile, for markers related to fibroblast, the expression as high in dermal fibroblast, but not hair follicle melanocytes (Figure 3l).

### UVB irradiation induces melanin synthesis by hair follicle melanocyte

UVB irradiation is known to be a double-edged sword that induces melanogenesis by epidermal melanocyte, while resulting in cellular senescence and cell death. On the other hand, it is generally regard that for certain cases, e.g., NB-UVB phototherapy for vitiligo, hair follicle melanocyte will be activated and migrate to epidermis (Nishimura et al., 2010; Quan et al., 2004), which results in perifollicular repigmentation. To investigate the effect of UVB irradiation on hair follicle melanocyte, we have measured relative melanin synthesis and DCT expression of hair follicle melanocyte. Both melanin synthesis and DCT expression has significantly increased after 6 days after UVB irradiation (Figure 4).

**Figure 4.**
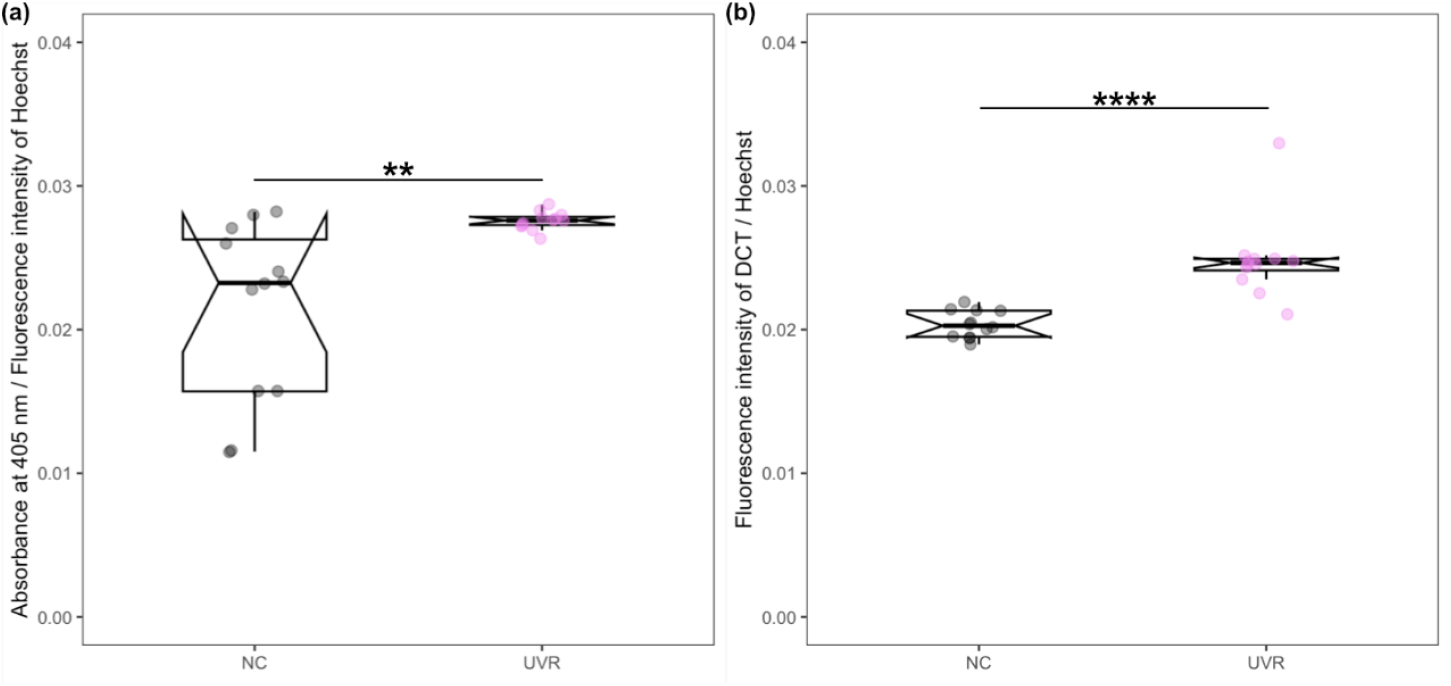
Melanin synthesis by hair follicle melanocytes after UV irradiation. (a) Relative melanin synthesis after UV irradiation. (b) Relative DCT expression of hair follicle melanocytes after UVB irradiation. Each plot represents 6 biological replicates and 2 technical replicates. Statistical significance were evaluated by Student’s t-test. * *p* < 0.05, ** *p* < 0.01, *** *p* < 0.001, **** *p* < 0.0001.

To understand the changes in hair follicle melanocyte which results in increased melanin synthesis, transcriptomic analysis of hair follicle melanocyte 21 h after UVB irradiation was conducted. tSNE plot has shown the changes in hair follicle melanocyte after UVB irradiation (Figure 5a). Comparing UVB irradiated hair follicle melanocyte to hair follicle melanocyte without UVB irradiation, 158 up-regulated genes and 8 down-regulated genes were identified (Figure 5b).

**Figure 5.**
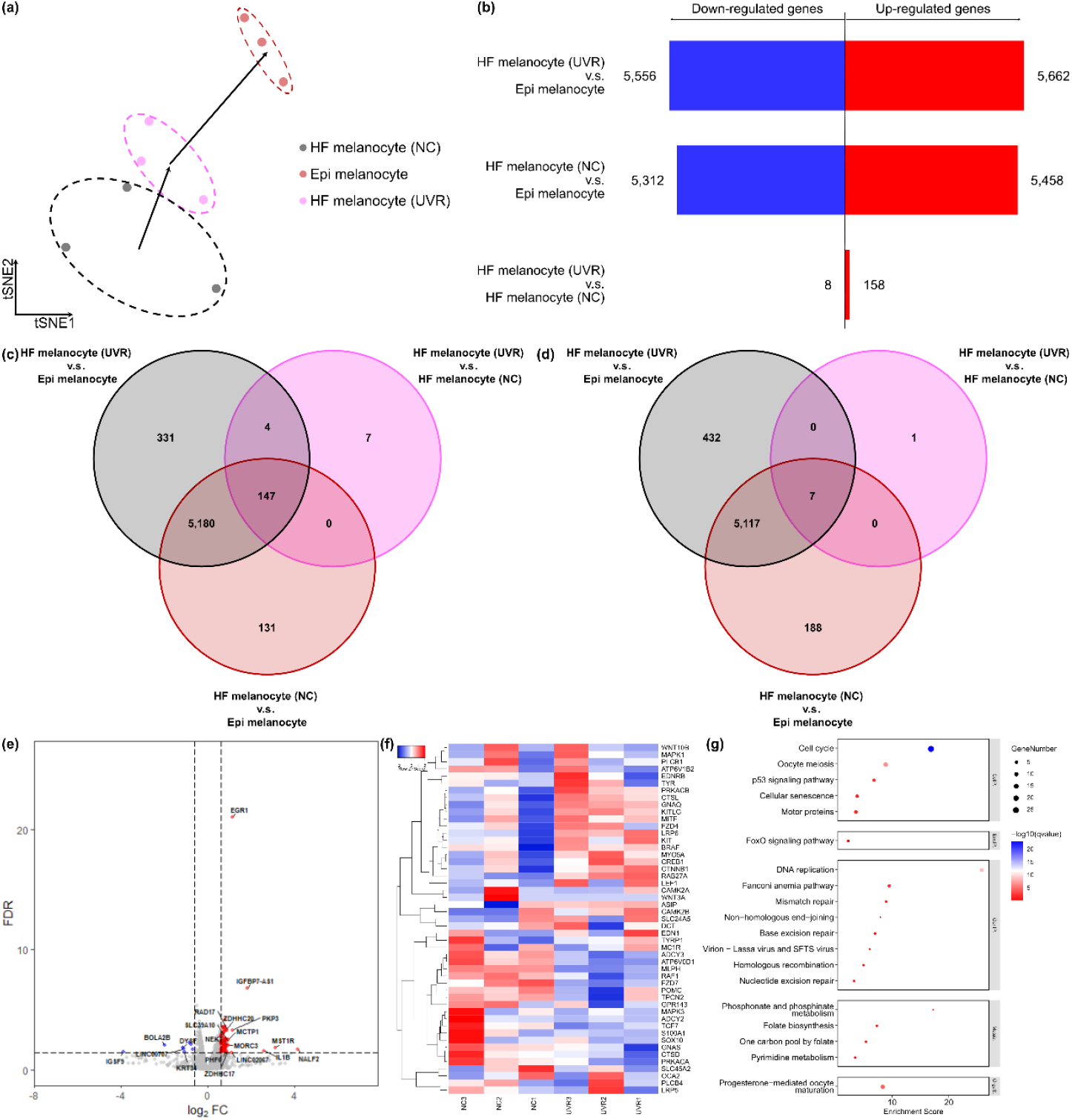
Transcriptomic analysis of UVB irradiated hair follicle melanocyte. (a) t-distributed Stochastic Neighbour Embedding mapping of hair follicle melanocyte, UVB irradiated melanocyte and epidermal melanocyte. (b) Plot of the number of up-regulated genes and down-regulated genes in each pair of samples investigated. Venn diagram of (c) up-regulated genes and (d) down-regulated genes of among each pair of samples investigated. (d) Volcano plot indicating the expression profile changes in hair follicle melanocyte after UVB irradiation. (f) Heatmap of genes related to melanogenesis and melanocyte differentiation. (g) Pathway enrichment of genes based on up-regulated genes obtained from hair follicle melanocyte after UVB irradiation.

151 up-regulated genes and 7 down-regulated genes were shared with those identified by comparing UVB irradiated hair follicle melanocyte to epidermal melanocyte (Figure 5c-d). Genes related to survival and proliferation of melanocytes (Figure 5e) along with p53 signalling pathway (Figure 5g) were up-regulated. Regulator of cell cycle arrest e.g., MST1R, p53 and CDKN1A were down-regulated, which suggested active cell proliferation. On the other hand, genes related to melanogenesis were also up-regulated (Figure 5f). This result suggested that UVB irradiation has induced cellular proliferation and melanogenesis by hair follicle melanocyte, which not only in align with observation in Figure 4, but also clinical observations.

### UVB irradiation induces migration and maturation of hair follicle melanocyte

To validate if UVB irradiation induces migration and maturation of hair follicle melanocytes as suggested in transcriptomic analysis, hair follicles were irradiated with 0.04 J cm^-2^ UVB and cultured for 4 days. The hair follicles were observed under microscope (Figure 6a). After UVB irradiation, the proximity of interfollicular epidermis and hair bulb were darken (Figure 6f – i) compared to control (Figure 6b – 6e), suggesting that melanocytes in hair follicle were activated, migrated into epidermis and hair bulb, then synthesise melanin.

**Figure 6.**
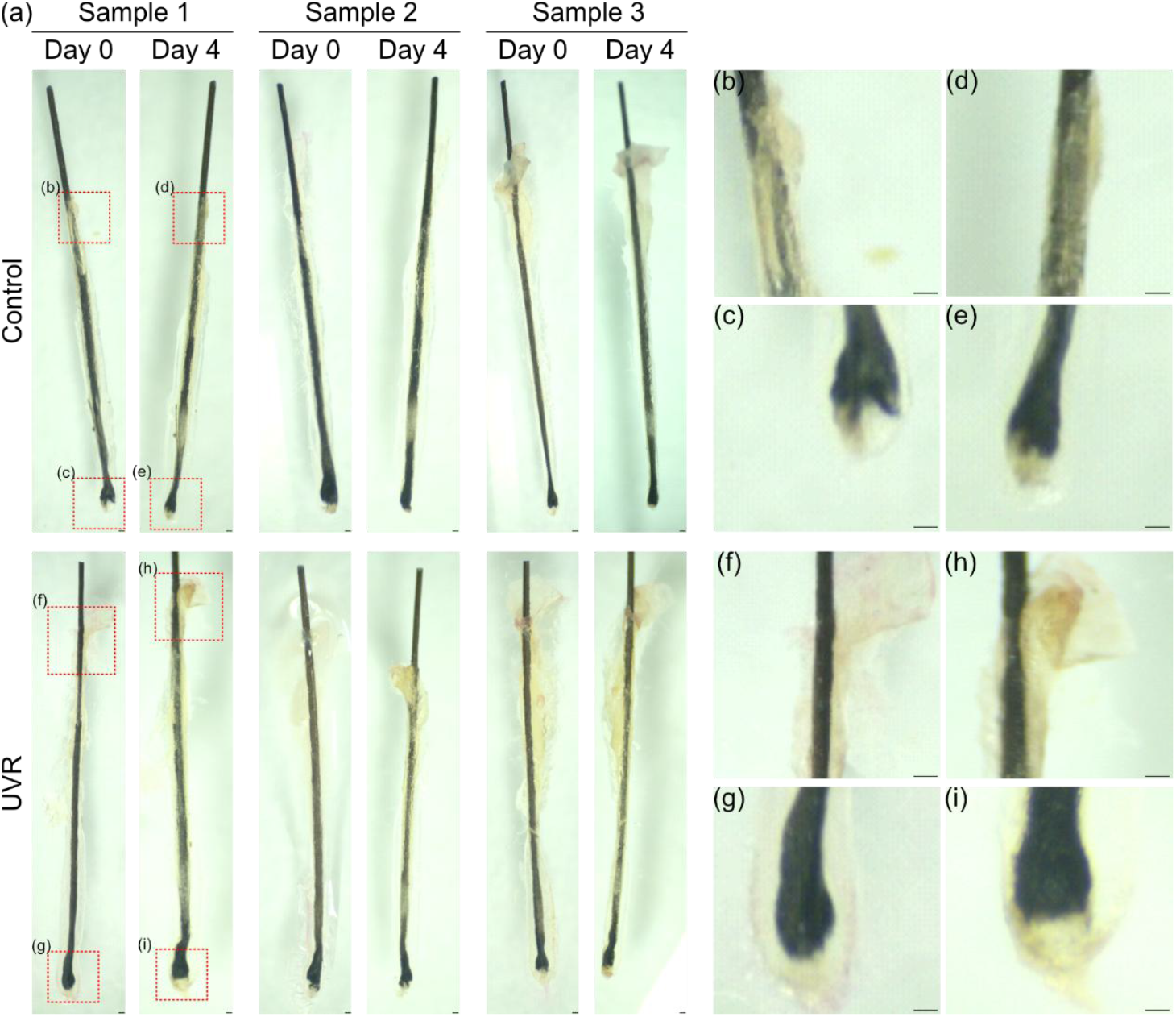
Migration and melanin synthesis by hair follicle melanocytes after UV irradiation. (a) Morphology of hair follicle immediately after UVB irradiation and cultured for 4 days after UVB irradiation. Control samples were hair follicle without UVB irradiation. Each set of data contains 3 hair follicles. Example of enlarged region of hair follicle at the proximity of interfollicular epidermis of (b, d) control and (f, h) UVB irradiated samples. Example of enlarged hair bulb of (c, e) control and (g, i) UVB irradiated samples. The scale bars are 200 μm.

## Discussion

Prior studies have isolated melanocytes from hair follicle and epidermis, and their morphological and antigenic differences have been identified. Amelanotic melanocyte from hair follicle and poly-dendritic melanocyte from epidermis are proliferative, while intensely pigmented hair follicle melanocyte is non-proliferative (Tobin & Bystryn, 1996). These melanocytes express different antigen on their surface, which the difference attribute to preferential destruction of hair follicle melanocyte in alopecia aerate, and epidermal melanocyte in vitiligo. While some other studies have described migration and differentiation of hair follicle melanocyte into epidermal melanocyte (Nishimura et al., 2010; Quan et al., 2004), the phenomenon has not been observed in human. To date, the characteristics of hair follicle melanocyte remain largely unknown. In this study, we have characterised hair follicle melanocyte in more detail, compare hair follicle melanocyte to epidermal melanocyte, and revealed the response of hair follicle melanocyte against UVB irradiation.

Through scRNA-seq analysis, we have identified a cluster of amelanotic hair follicle melanocyte, which express high level of neural stem cells markers, e.g., PAX3, SOX5, and nestin. This subpopulation of hair follicle melanocyte is potentially melanocyte stem cell which retain markers from neural crest stem cell (Mort et al., 2015), and they can differentiate into melanocytes in interfollicular epidermis, followed by epidermal melanocyte.

Then, hair follicle melanocytes were isolated and compared to epidermal melanocyte and dermal fibroblast. While hair follicle melanocytes are NKI/beteb^+^ cells as described in previous studies, they also express MITF. Interestingly, this subpopulation of melanocyte has high CD90 expression, which is a stem cell marker, suggesting the stem cell property of hair follicle melanocyte.

Analysing the transcriptomic profile of hair follicle melanocyte shows that hair follicle melanocytes are less differentiated pigment producing cells.

Prior study has suggested that epidermal melanocytes undergo cell death after UVB irradiation (Xu et al., 2025). We have also found that UVB irradiation, particularly above 0.04 J cm^-2^ will result in retarded proliferation and cell death (Figure S1). Unlike epidermal melanocyte which undergo cell death after UVB irradiation, hair follicle melanocyte proliferated, synthesised DCT, followed by melanin. This is evidenced by the upregulation of DNA replication, cell replication and cell cycle related genes at 21 h after UVB irradiation, and higher DCT and melanin concentration 6 days after UVB irradiation. In vitro cell proliferation assay has also suggested that up to 0.3 J cm^-2^ UVB irradiation did not induce changes in proliferative capability of hair follicle melanocytes (Figure S2). This supports the inference that hair follicle melanocyte proliferates, migrate to epidermis and produce melanin after UVB irradiation, which might result in skin repigmentation.

In in situ study based on hair follicles, we found that the root sheaths of hair follicles darken after UVB irradiation (Figure 6), suggesting UVB might induce migration and maturation of melanocytes in hair follicle, which differ from the observation based on epidermal melanocytes. As the melanocytes have also migrated upward, this suggests the potential that hair follicle melanocytes might serve to repigment epidermis, particularly acting in repigmentation of vitiligo lesions after receiving UVB phototherapy.

In summary, the results presented in this study give important insights into the property of melanocyte in human hair follicle, their molecular characteristics, and their response against UVB irradiation which contribute to skin pigmentation. We could distinguish between epidermal melanocyte and hair follicle melanocyte in human. Importantly, discovery of the proliferation of hair follicle melanocyte after UVB irradiation is essential for understanding how phototherapy is useful in skin pigmentation disease.

## Materials and Methods

### Ethical approval

This study uses leftover samples after surgical procedures. This study was approved by the Ethics Committee of Huashan Hospital of Fudan University (KY2020-793, KY2020-565).

### Materials

Levo-dihydroxyphenylalanine was obtained from Sigma (People Republic of China), Goat antibody against human microphthalmia-associated transcription factor (MITF), NorthernLights™ NL557-conjugated donkey antibody against goat immunoglobulin G (IgG), NorthernLights™ NL493-conjugated donkey antibody against mouse IgG, NorthernLights™ NL557-conjugated goat antibody against mouse IgM, sheep antibody against human cluster of differentiation (CD) 90, NorthernLights™ NL557-conjugated donkey antibody against sheep IgG were obtained from R&D Systems (Minneapolis, United States of America), mouse antibody against human NKI/beteb was obtained from Abcam (Cambridge, United Kingdom), mouse antibody against human fibroblast surface protein (FSP) was obtained from Novus Biologicals (Colorado, United States of America), FITC-conjugated mouse antibody against human dopachrome tautomerase (DCT) was obtained from Santa Cruz Biotechnology (Santa Cruz, United States of America).

### Single cell transcriptomic analysis of melanocytes

Single cell RNA sequencing data from normal human skin was obtained from public dataset China National Center for Bioinformation: SRX14231218) and dataset on hair follicle was sequenced (in preparation for publication). Seurat package (v5.3.0) and monocle3 (v1.4.26) were used for bioinformatics data analysis (Hao et al., 2024; Qiu et al., 2017).

### Isolation of epidermal melanocytes

Epidermal melanocytes were isolated with protocol modified from Goff et al. (2023). Briefly, human skin was sterilised and treated with 10 mg ml^-1^ dispase II overnight at 4°C. Then, epidermis was peeled from dermis, and the epidermis were placed in 0.25% trypsin-ethylenediaminetetraacetic acid solution and incubated for 60 mins. The epidermis was dispersed and the reaction was inhibited by supplementation of equal volume of Dulbecco’s modified Eagle medium containing foetal bovine serum (FBS). The remaining tissue was removed by filtration through 40 μm cell strainer, and the filtrate were pelleted by centrifugation at 800 xg for 5 mins, followed by resuspension in epidermal melanocyte culture medium (medium 254 supplemented with human melanocyte growth supplement-2). The number of cells was counted with haemocytometer, and approximately 4 × 10^4^ cells cm^-2^ were plated on type IV collagen coated cell culture dish. For the first 4 days of cultivation upon isolation, 50 μg ml^-1^ geneticin was supplemented to culture medium to eliminate potential contamination by mesenchymal cells.

### Isolation of hair follicle melanocytes

Hair follicle melanocytes were isolated from hair follicles by trypsin digestion. After sterilisation, hair follicles were incubated in 0.25% trypsin-ethylenediaminetetraacetic acid solution for 10 mins. Hair follicles were dispersed, and the reaction was inhibited by supplementation of equal volume of Dulbecco’s modified Eagle medium containing FBS. Hair shafts were removed by filtration through through 40 μm cell strainer, and the filtrate were pelleted by centrifugation at 800 xg for 5 mins, followed by resuspension in hair follicle melanocyte culture medium (proprietary medium; can be purchased from ReMed Regenerative Medicine Clinical Application Institute). The number of cells was counted with haemocytometer, and approximately 4 × 10^4^ cells cm^-2^ were plated on type IV collagen coated cell culture dish. For the first 4 days of cultivation upon isolation, 50 μg ml^-1^ geneticin was supplemented to culture medium to eliminate potential contamination by mesenchymal cells.

### Isolation of dermal fibroblast

Dermis collected after separation of epidermis for melanocytes isolation were diced and placed in type IV collagen-coated cell culture dish. Then, add DMEM containing 10% foetal bovine serum and 1% antibiotic-antimycotic into the dish. Medium was replaced every 3 days until sprouting of fibroblast was observed.

### Levo-dihydroxyphenylalanine staining

After passaging, cells were seeded on poly-*L*-lysine-coated slides and cultured to approximately 60% - 80% confluency. The slides were retrieved, rinsed with Dulbecco’s phosphate buffer saline solution (D-PBS), and the cells were fixed with 4% paraformaldehyde for 30 min. The cells were rinsed twice with D-PBS and incubated with 0.1% Levo-dihydroxyphenylalanine (*L*-DOPA) solution at 37°C for 4 h. *L*-DOPA solution was removed, and the nuclei were counter-stained with haematoxylin. Stained cells were observed under brightfield microscope.

### Immunocytochemistry

The cells cultured on poly-*L*-lysine-coated slides were rinsed with D-PBS and fixed with 4% paraformaldehyde for 30 mins at room temperature and pressure. Blocking, incubation with primary and secondary antibody were conducted described by manufacturer’s instructions. No primary control was prepared by incubating with D-PBS instead of primary antibody. Hoechst 33258 were used as counterstain. The cells were washed with D-PBS and micrographs were obtained using fluorescence microscope (IX73, Olympus, Tokyo, Japan) with no primary control used as background correction.

### Transcriptomic analysis of hair follicle melanocytes

After cultivation of dermal fibroblast, epidermal melanocyte, and hair follicle melanocyte to 80% confluency, total RNA was isolated with TRIeasy® total RNA extraction reagent (Yeasen Biotechnology, 10606ES60, Shanghai, People Republic of China). Then, the total RNA extracts were snap-frozen at -80°C until further processing. Before library construction, the purity of RNA was analysed with microphotospectrometer (NanoDrop 2000, Thermo Scientific, USA) and the integrity was analysed with a bioanalyzer (Agilent 2100 Bioanalyzer, Agilent Technologies, California, USA). Then the libraries were constructed using VAHTS Universal V10 RNA-seq Library Prep Kit (Premixed Version) according to the manufacturer’s instructions. Transcriptome sequencing was purified and sequenced by Shanghai OE Biotech Co. Ltd. After construction of libraries, the products were sequenced on an Illumina Novaseq X Plus platform. 150 bp paired-end reads were generated. About 31.23 million raw reads were generated for each sample. Raw reads in FASTQ format were processed using fastp (Chen et al., 2018), and low-quality reads were removed to obtain clean reads. The clean reads were mapped to reference genome (GRch38.p13) using HISAT2 (Kim et al., 2015). Further analyses were conducted using R (v4.5.1).

### UVB irradiation of hair follicles melanocytes

After cultivation of hair follicles melanocytes to 80% confluency, the cells were collected by trypsinisation and pelleted by centrifugation. Then, the cells were resuspended with D-PBS to 0.5 × 10^7^ cells ml^-1^ density. 0.1 ml cell suspension were seeded into 3 wells in each well of 6-well plate, to form a thin film of cell suspension. Then, the 6-well plate was exposed to 0.04 J cm^-2^ 312 nm UV (Scientz03-II, Ningbo Scientz Biotechnology Co., Ltd., Ningbo, China). Then, the 3 wells were filled up with hair follicle melanocyte culture medium to a final volume of 3 ml. The remaining 3 wells were seeded with 0.5 × 10^6^ cells in 3 ml hair follicle melanocyte culture medium, which will be used as control. The number of cells were adjusted according to growth area when 24-well plate was used.

To evaluate relative melanin synthesis capability of hair follicle melanocytes after UVB irradiation, the cells were irradiated with UVB, cultured with hair follicle melanocyte culture medium in 37°C, 5.0% CO2 incubator, and the medium was replaced every other day. The cells were retrieved at 6 days after UVB irradiation. The cells were rinsed and fixed with paraformaldehyde, stained with Hoechst 33258, and immerse in D-PBS. Then, absorbance at 405 nm and fluorescence intensity at 352/461 nm was measured.

For bulk RNA-seq, total RNA was isolated from melanocytes 21 h after UVB irradiation with TRIeasy® total RNA extraction reagent. Then, the total RNA extracts were snap-frozen at -80°C until further processing. Transcriptomic analysis was conducted as described in previous section.

### In situ study based on hair follicle

After removal adipose tissue and dermis attached to hair follicles, the hair follicles were rinsed with D-PBS. Then, hair follicles were irradiated with 0.04 J cm^-2^ 312 nm UV, then transferred into hair follicle culture medium (proprietary medium; can be purchased from ReMed Regenerative Medicine Clinical Application Institute). The hair follicle culture medium was replaced every other day. The hair follicles were observed with stereomicroscope (SMZ745T, Nikon, Tokyo, Japan) at day 4 after cultivation.

## Acknowledgement

Not applicable

