## Supplementary material for "Isolation and Characterisation of Hair Follicle-Derived Melanocytes": Figure S1; Figure S2

Wenyu Wu (Academic)

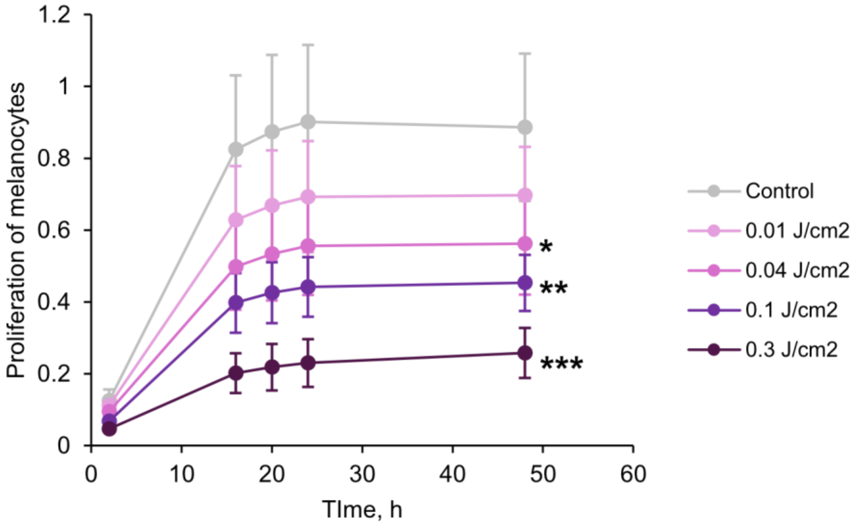


**Figure S1. Effect of UVB on epidermal melanocytes.** Each plot represents 5 biological replicates. Statistical significance of each group against control was evaluated by Student's t-test. * $p < 0.05$, ** $p < 0.01$, *** $p < 0.001$.

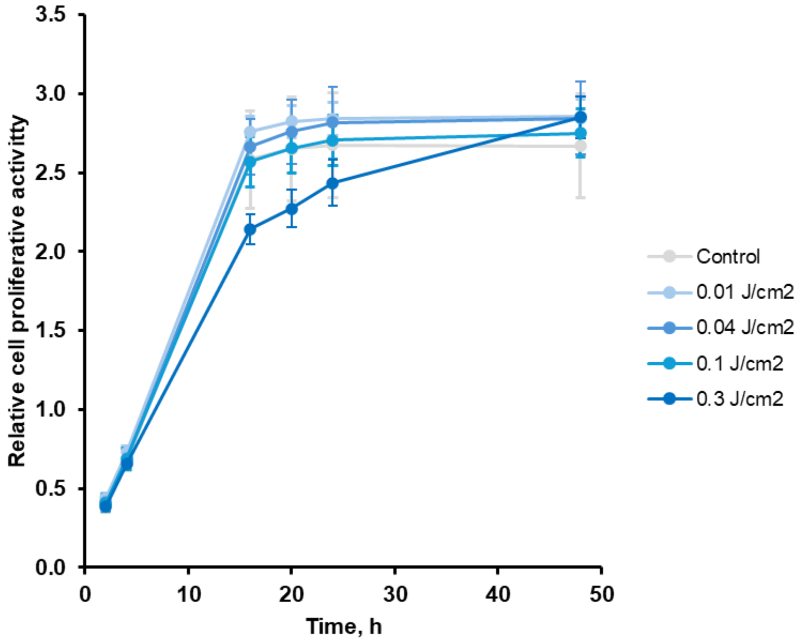


**Figure S2. Effect of UVB on hair follicle melanocytes.** Each plots represents 6 biological replicates. Statistical significance of each group against control was evaluated by Student's t-test.
